# Modeling time-series data from microbial communities

**DOI:** 10.1101/071449

**Authors:** Benjamin J Ridenhour, Sarah L Brooker, Janet E Williams, James T Van Leuven, Aaron W Miller, M Denise Dearing, Christopher H Remien

## Abstract

As sequencing technologies have advanced, the amount of information regarding the composition of bacterial communities from various environments (e.g. skin, soil) has grown exponentially. To date, most work has focused on cataloging taxa present in samples and determining whether the distribution of taxa shifts with exogenous covariates. However, important questions regarding how taxa interact with each other and their environment remain open, thus preventing in-depth ecological understanding of microbiomes. Time-series data from 16S rDNA amplicon sequencing are becoming more common within microbial ecology, but given the ‘big data’ nature of these studies, there are currently no methods capable of utilizing the breadth of the data to infer ecological interactions from these longitudinal data. We address this gap by presenting a method of analysis using Poisson regression fit with an elastic-net penalty that 1) takes advantage of the fact that the data are time series; 2) constrains estimates to allow for the possibility of many more interactions than data; and 3) is scalable enough to handle data consisting of thousands of taxa. We test the method on gut microbiome data from white-throated woodrats (*Neotoma albigula*) that were fed varying amounts of the plant secondary compound oxalate over a period of 22 days to estimate interactions between OTUs and their environment.

## Introduction

Methodological advances in DNA sequencing have uncovered microbial diversity that extends far beyond that which could be detected using traditional cell culture methods. Because of the ease and inexpensive nature of new technologies, researchers are collecting increasing amounts of data with regard to various microbiomes (e.g. skin, soil, gut), a trend which will only increase with newly created funding sources such as the recently announced U.S. National Microbiome Initiative (The White House Office of Science and Technology Policy, 2016). To date, most studies center around identifying members of the community using 16S rDNA sequencing and using diversity measures and ordination techniques to compare samples (Ramette, 2007; Cassman *et al.*, 2016). While such analyses yield a large amount of information regarding where and when a particular microbe might be found, they tell almost nothing about why the microbe is there, how it interacts with its environment (e.g. other microbes, host), and what functions it may be providing toward—or detracting from—the overall ecosystem-level services.

To address some of these central questions of interactions and function, ecological and evolutionary theory developed for macro systems has begun to be applied to microbial systems. For example, microbiome data derived from 16S rDNA sequencing has been used to estimate population dynamics of microbial communities (Marino *et al.*, 2014), to infer how communities respond to perturbations (Stein *et al.*, 2013), and to assess important community properties such as stability and resilience (Coyte *et al.*, 2015). Many of these ideas that are central to microbial ecology and microbiome function are inherently dynamic, and as such require longitudinal data.

Unfortunately, the staggering number of operational taxonomic units (OTUs) present in microbiomes prevents straightforward application of traditional ecological modeling methods, so methods to analyze the data have lagged behind collection. Whereas a large macro ecological system may track up to one hundred species (e.g. Montoya & Solé, 2002), microbial communities often have thousands of OTUs, which provides a significant hurdle for estimating the key interactions between microbes and other microbes and their environment. Various simplifications of data and models have been used to deal with this issue; the most common of which are to vastly reduce the size of the data by either aggregating the data at certain taxonomic levels (e.g. treating Alphaproteobacteria as a model factor; McGeachie *et al.*, 2016), sub-setting the data into a few taxa of interest because they are believed to be important (Hunt *et al.*, 2011), or both (Olesen *et al.*, 2016). Aggregation is a particularly specious practice because of heterogeneity within aggregated taxa. For example, estimating how Alphaproteobacteria interact with Gammaproteobacteria is akin to estimating how all dicotyledonous plants interact with monocotyledonous plants. Similarly, while focusing on only a few taxa of interest can make statistical inference techniques tractable, the interactions that may actually be driving the dynamics may be left out of the model. Ecologically important forces, such as trait-mediated indirect interactions (Ridenhour & Nuismer, 2012; Berry & Widder, 2014), will be missed in this type of analysis.

Another recent method of analyzing microbiome data to infer drivers of ecological dynamics is to compare large-scale patterns. For example, Bashan *et al.* (2016) used patterns of dissimilarity and overlap between microbiomes to infer whether interactions within a microbial community were host driven or not. However, using an analysis like this ignores several ecological principles, such as nutrient flow in ecosystems (Jordano *et al.*, 2003), that are likely relevant in microbial systems. For example, some bacteria must be present within the community that can convert host resources (e.g. glycogen) into resources that the microbial community can subsequently utilize; i.e. primary producers must be present that interact with the host.

Ideally, we desire ecologically relevant methods that are capable of utilizing all information gathered from sequencing to robustly infer relationships between OTUs and their environment. Methods for model estimation using big data where the number of possible explanatory variables is larger than the number of observations (*p* ≫ *n*) typically involve the use of regularization (Tibshirani, 1996; Zou & Hastie, 2005; Meinshausen & Bühlmann, 2010) to eliminate potential explanatory variables and infer robust, stable predictive models. These regularization techniques have been successfully applied in many fields where big data are common, such as gene expression data and proteomics (Xing *et al.*, 2001). Furthermore, related techniques have been applied to microbiome research. For example, Kurtz *et al.* (2015) used a form of a graphical lasso procedure (sparse inverse covariance estimation) (Friedman *et al.*, 2008) applied to relative abundance data for entire microbiome samples. Their study demonstrates that regularization techniques can successfully be applied to 16S sequencing data at the scale of big data, allowing for downstream statistical inference. Using regularization methods can avoid misleading interpretations caused by aggregating data or arbitrarily studying certain species within a microbial community.

Here, we present a novel method of analyzing 16S sequencing data that utilizes untransformed count data from the entire community and relies on regularization to infer interactions. We focus on applying this method to time-series data, which is a rapidly expanding microbiome research area and an area of special need for such techniques. We emphasize however that the methods presented here are not limited to the analysis of time series and are broadly applicable to related microbiome analyses. As an example of the power of this technique, we apply the method to gut microbiome data collected from *Neotoma albigula* (white-throated woodrats) during an ∼3 week feeding trial in which the subjects were fed oxalate, a plant secondary defensive compound.

## Methods and Subjects

### Modeling Strategy

We used an ARIMA model with Poisson errors fit with elastic-net regularization to estimate robust predictive models of microbiome dynamics. ARIMA models are commonly used in the analysis of time-series data because they provide a flexible framework that can accommodate many autocorrelation structures, stationarity conditions, and seasonality (Ives *et al.*, 2003). The choice of Poisson distributed errors is critical to avoid issues related to compositional data: raw read counts and total read counts are the data analyzed rather than transformed compositional data. The Poisson distribution is a natural choice for count data (Anders & Huber, 2010), and, furthermore, by using the total read count as the offset in a log-linked Poisson regression model, the zeroes observed in the data are treated appropriately and have consistent meaning across variable total read counts. The resulting full model is

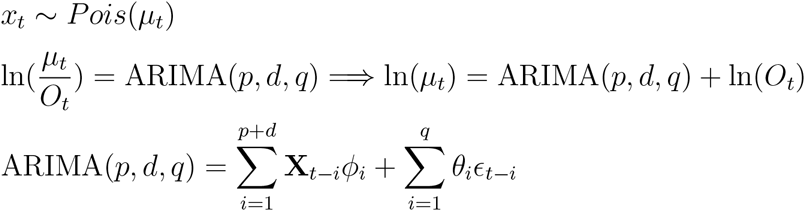

where subscripts indicate the time of observation, *x* is the number reads observed for a particular OTU, *µ* is its mean, *O* is the total number of reads, **X** is the vector of predictor variables (i.e. other OTUs, covariates, etc), ε is the residual error, and *ϕ* and *θ* are the estimated model parameters (though we are principally interested in *ϕ* because this vector contains the estimated interactions between OTUs). Note that the number of parameters in the full model is (*p* + *d*)|**X**| + *q*, thus increasing either *p* or *d* can have large effects on the number of parameters estimated when the number of predictor variables (|**X**|) is large.

The full model represents a flexible way to model interactions between species that takes full advantage of the data type and its time-series structure, but would be highly overparameterized for nearly all microbial community data because of the large number of predictor OTUs. It is commonly held that most species interact with few other species (Faust & Raes, 2012) and some propose exponential or scale-free distributions to the number of edges in interactions networks (Fernandez *et al.*, 2015; Kurtz *et al.*, 2015). Regardless, the ecological expectation is that a fully saturated model, such as the one above, is not realistic. We therefore employ a regularization algorithm to select robust interaction models that have a minimal number of parameters. The elastic-net regularization is a highly flexible and rapid algorithm that uses both the 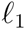 and 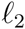 norms (i.e. lasso and ridge regression respectively) (Tibshirani, 1996; Draper & Pukelsheim, 2002; Zou & Hastie, 2005). To estimate the parameters 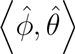 elastic-net algorithm solves the equation

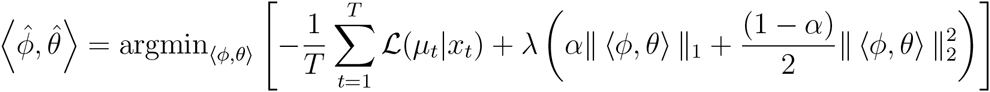

where 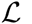 is the log-likelihood of the observed data (*x*_t_) given the modeled mean (*µ*_t_), 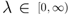 controls the strength of the elastic-net penalty (λ = 0 is equivalent to standard leastsquares regression), and 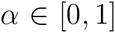 blends the penalty due to the 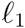 and 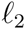 norms (*α* = 0 is ridge regression and *α* = 1 is lasso regression) (Tibshirani, 1996). Cross-validation techniques are used to choose optimal values for these parameters. The use of the elastic-net approach in combination with a Poisson ARIMA model allows our method to filter through large numbers of OTUs and robustly model changes in a microbiome over time.

### Application of Model to Oxalate Degradation in *Neotoma albigula*

We used our modeling strategy to estimate microbial community dynamics from 16S rDNA time-series data collected from the white-throated woodrat, *Neotoma albigula* (Miller *et al.*, 2016). These animals were experimentally fed varying amounts of oxalate, a naturally occurring plant secondary compound that has been demonstrated to have toxic effects on a broad range of herbivores (e.g. insects, mammals) (Allison *et al.*, 1985; Dearing *et al.*, 2005). Plants create crystalline structures known as raphides when a surplus of calcium oxalate is present; these crystals are needle-shaped and physically damage the intestinal tract of herbivores. This physical damage may also facilitate delivery of other toxins (e.g. proteases) through the wall of the digestive tract (Miller *et al.*, 2000; Franceschi & Nakata, 2005). Direct mortality, decay of the mouth and gastrointestinal tract, gastric hemorrhaging, and diarrhea have all been observed in mammals that consume large quantities of oxalate (Miller *et al.*, 2014). Of human relevance, many kidney stones form from calcium oxalate, which may arise due to oxalate rich diets; the pain associated with passing kidney stones at least partly stems from similar needlelike structures (Miller *et al.*, 2000; Franceschi & Nakata, 2005).

*N. albigula* rely on cactus, particularly *Opuntia*, for their diet (Justice, 1985; Miller *et al.*, 2014), which are known to have high concentrations of calcium oxalate (Shirley & Schmidt-Nielsen, 1967). Mammals however are not known to have any mechanisms for metabolizing this toxic compound but are known to harbor bacteria capable of the task within the gut (Hodgkinson, 1977; Allison *et al.*, 1985; Turroni *et al.*, 2007). Prior studies of white-throated woodrats have shown that the microbiota of their gut has numerous oxalate-degrading taxa including—but not limited to—*Oxalobacter formigenes*, *Lactobacillus*, *Bifidobacterium*, *Streptococcus*, and *Enterococcus* (Allison *et al.*, 1985; Jones & Megarrity, 1986; Kageyama *et al.*, 1999; Hokama *et al.*, 2000; Sundset *et al.*, 2010). *O. formigenes* has been of particular interest within the gut community because it is known to require oxalate as a carbon and energy source (Allison *et al.*, 1985). It has been hypothesized that the specialization of and coevolution with the gut microbiome is the reason *N. albigula* may consume levels of oxalate that would be lethal for many other mammals and digest ≥90% of this defensive compound (Shirley & Schmidt-Nielsen, 1967; James & Butcher, 1972; Justice, 1985; Palgi *et al.*, 2008).

### Feeding Trials

We collected gut microbiome data from 6 wild-caught *N. albigula* trapped at Castle Valley, Utah in October 2012. Animals were transported back to the University of Utah Department of Biology Animal Facility and held in captivity for six months prior to experimentation. During this time, animals were fed a high-fiber rabbit chow (Teklad formula 2031; Harlan, Denver, CO, USA), which contained a baseline amount of oxalate.

Once the trial began, oxalate concentrations within the animals' food was incrementally increased for 17 days and then dropped to initial level for an additional five days. The amount of oxalate consumed and excreted was measured for each woodrat. Fecal pellets were collected and then underwent high-throughput 16S rDNA amplicon sequencing to determine the OTUs present in guts of the animals. OTU read counts from the cleaned and processed data were then analyzed using the model described above. A general overview of the workflow for the analyses is provided in Figure 1.

**Figure 1:**
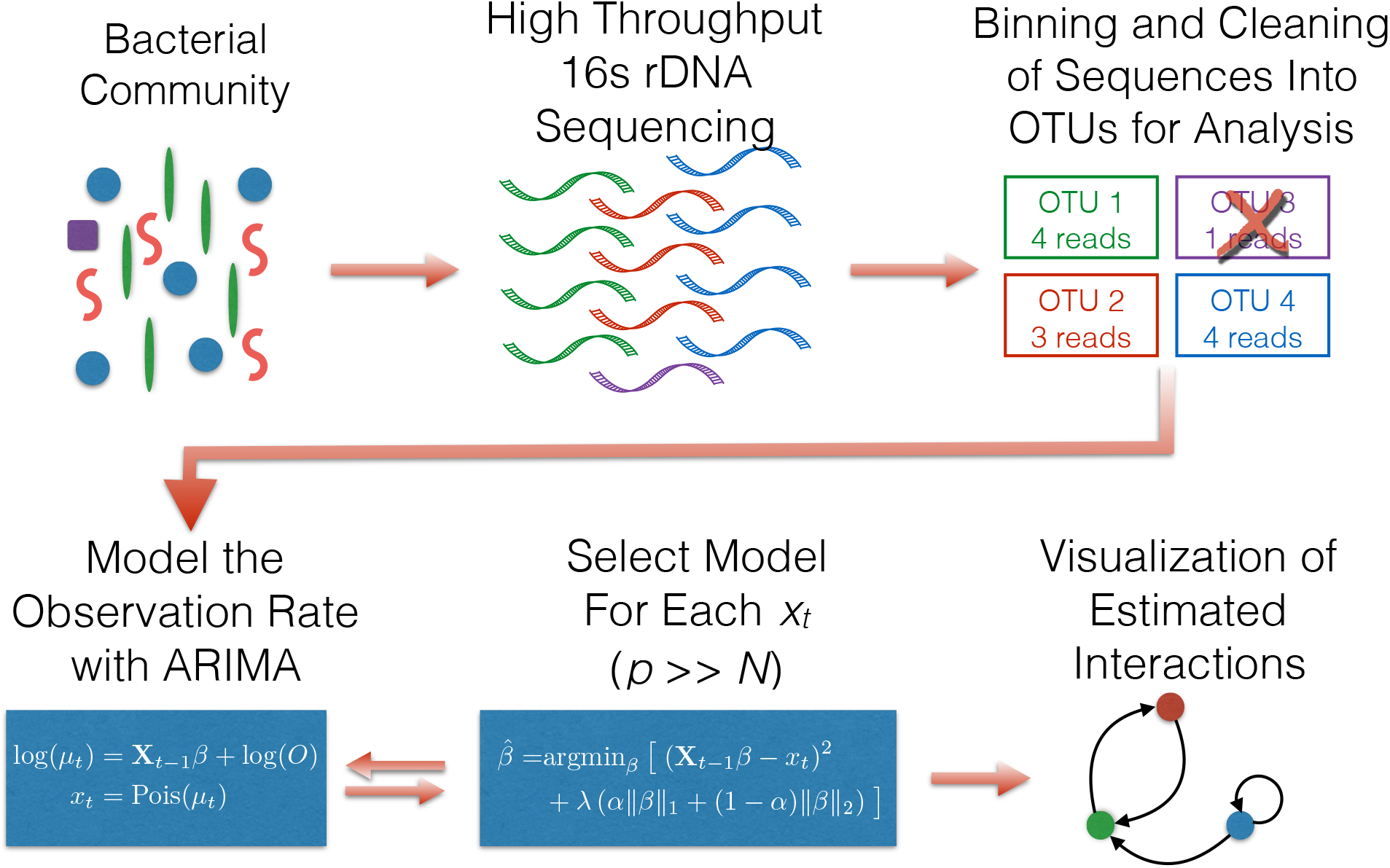
The general workow for analyzing 16S rDNA data using a regularized ARIMA model with Poisson errors. The first several steps are the typical sequencing and bioinformatic practices where sequences are obtained and cleaned using programs such as QIIME. An additional step of dropping particularly low read count OTUs may be necessary to avoid problems with the statistical analyses reporting errors. Afterward, the cleaned data are passed to the regularization algorithm to fit an appropriate ARIMA model. The final step is to analyze the estimated interaction network (e.g. heatmaps, networks, summary statistics) to interpret the models returned from the analysis.

To quantify the effect of oxalate on the gut microbiota, a custom 0.2% oxalate diet was formulated (Harlan, Denver, CO, USA) and mixed with high-fiber rabbit chow in a 3:1 ratio to give a 0.05% baseline oxalate feeding level. Additional concentrations of 0.5%, 1%, 1.5%, and 3% by dry weight were achieved by adding sodium oxalate (Fisher Scientific, Pittsburgh, PA, USA) directly to the diet. The oxalate diets were given to animals in sequence for three days each, with the exception of the 0.05% oxalate diet, which was given for five days both at the beginning and the end of the diet trial. This schedule produced observations on day 5 (*t*_0_), 8 (*t*_1_), 11 (*t*_2_), 14 (*t*_3_), 17 (*t*_4_), and 22 (*t*_5_). Food and water were given *ad libitum* in metabolic cages, which were used to separate and collect urine and feces from each individual animal. Oxalate consumed was quantified from food intake and oxalate concentration, while oxalate excreted was quantified from urine and feces. These metrics were used to quantify total oxalate degradation, which we defined as the difference between oxalate consumed and oxalate excreted.

To track changes to the gut microbiota, feces were collected from each animal on the last day of each dietary period, thus maximizing the effect of the specified oxalate concentration on the gut microbiota. Feces were collected from the top of the 50ml conical tube to ensure minimal exposure to aerobic conditions, and immediately frozen at -80C until DNA extraction. DNA extractions were performed with the QIAmp DNA stool minikit (Qiagen, Germantown, MD, USA). Microbial inventories were generated by amplifying the V4 region of the 16S rDNA gene with primers 515F and 806R (Caporaso *et al.*, 2012) on an Illumina MiSeq at Argonne National Laboratory (Chicago, IL, USA).

### Data Processing

Sequence data were processed and demultiplexed in QIIME (Caporaso *et al.*, 2010) using the default quality control parameters. Sequences were binned into OTUs with a de novo picking strategy using UCLUST (Edgar, 2010) at a minimum sequence identity of 97%. Chimeras were removed with ChimeraSlayer (Haas *et al.*, 2011) along with sequences identified as chloroplasts or mitochondria.

### Computational Details

All statistical analyses were performed using R v3.2.2 (R Core Team, 2014) with glmnet v2.0-2 (Friedman *et al.*, 2010), and all R scripts are available upon request. Prior to beginning the analyses, we eliminated any OTUs in the data for which there were a small number (< 6) of average reads per sample because they lacked sufficient data for statistical analysis. Eliminating these unanalyzable OTUs resulted in 90% reduction in the number of OTUs giving a microbial community that had 624 OTUs for modeling. The rationale for this cleaning is that there must be enough data for an OTU to successfully run a statistical model; if there is too little variation among samples for an OTU, then the statistical modeling will fail as there is no information.

The glmnet function in the glmnet package has a number of options for performing model regularization. The most important parameters are λ and *α* which control the penalization. To find an optimal combination of these parameters, we used the built in cross-validation function provided in the glmnet package (“cv.glmnet”) to loop across various levels of λ (100 values by default). We simultaneously looped across levels of *α* ranging from 0.5 to 1.0 in steps of 0.1. Because the cross-validation step performs K-fold cross-validation, the data folds are random; we therefore ran 500 replicates per *α* level to get the average cross-validated deviance for a particular *α*, λ combination. The best model was chosen for each *α* level, and the final model was chosen by utilizing AIC values and comparing between those best models. Other methods exist for choosing this parameter combination (see the c060 R package by Sill *et al.*, 2014, for an example of another method), but testing various algorithms is beyond the scope of this article. Other than the choice of λ and α, default parameter settings were passed to glmnet, with the exception of the “grouped = FALSE” argument to ensure enough observations per fold in the default 10-fold cross-validation scheme.

We performed two different variants of the model. The first of these was a model in which oxalate consumption was forced to be a variable within the model (i.e. oxalate consumption was a parameter that was not subject to the regularization penalties). The justification behind forcing this variable is that the experiment was designed to detect the influence of oxalate on the gut microbiome of the subjects. For comparison purposes, a second model was run where oxalate consumption was part of the regularization scheme and thus could either remain in, or be left out of, the final model chosen by the glmnet algorithm. These two models represent common research scenarios: determining effects of a particular factor that was experimentally manipulated and “natural” experiments where potential covariates change in an uncontrolled manner.

Post-analysis cleaning of models for interpretation purposes was minimal. Models having a pseudo-*R*^2^ < 0.02 were discarded from the results; doing so eliminated models for 38 OTUs in our data and left 580 models for interpretation. Effect sizes were determined by multiplying the mean OTU read counts by the corresponding estimated parameter 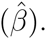 Estimated observation rates were calculated using the predict function in R. The Greengenes bacterial taxonomy database `“gg_13_5_taxonomy”` was searched to identify the phylogenetic relatedness of taxa identified in this study. The R package ape v3.5 (Paradis *et al.*, 2004) was used to parse and plot the reduced Greengenes phylogeny.

## Results

The experimental setup allowed us to examine how the gut microbiome of six woodrats changed over a three week period with varying levels of oxalate consumption. We wished to infer both how OTUs within the microbial community interacted with each other, as well as to estimate how OTUs were affected by oxalate concentration. To do so, we used an ARIMA(1,0,0) (i.e. AR(1)) model structure to limit the complexity of the model given the limited number of samples. We also included exogenous covariates for the amount of oxalate consumed and subject effects in the design matrix, bringing the total number of potential explanatory variables to 631.

We will focus on the results of the model where oxalate consumption was forced into the AR(1) model and highlight differences between it and the “unforced” model. The use of elastic-net regularization easily accommodated our analysis which included 624 OTUs and 7 other potential covariates (in other trials—not shown here—the method has worked for data sets containing thousands of covariates). We applied the AR(1) model to 626 dependent variables (624 OTUs, oxalate digested, and oxalate excreted); of these, analyses of 8 OTUs failed due to their patterns of presence/absence (all 8 were only observed in one woodrat and in only 2 of that subject's six samples). Of 618 × 631 = 389 958 potential parameters, the elastic-net algorithm selected models that had a total of 4174 parameters (∼ 1%); the unforced model produced 3556 parameters in comparison. Of the 618 dependent variables, 489 had a model that included additional variables from the minimal model that consists of only an intercept and oxalate consumed (469 for the unforced model). Thus, the regularization procedure produced models where there were relatively few predicted interactions between OTUs within the gut microbiome (i.e. a sparse interaction matrix).

The estimated network of interactions between species fits the “small-world” network paradigm that has been observed in many other non-microbial communities (Watts & Strogatz, 1998; Montoya & Solé, 2002). Figure 2 shows the in- and out-degree distributions for the predicted interaction network (i.e. the number of covariates affecting and affected by an OTU, respectively). These distributions fall in between what would be expected in a scale-free network and a random network. The average path length in the estimated network is *L* = 0.130, whereas a similar random network would have an average path length of *L_random_* ≈ 0.077. The average clustering coefficient (transitivity) of the estimated interaction network is *C* = 0:357 which compares to *C_random_* ≈ 0.020 in the random network. Therefore, our network fits the definition of a small world where *L*_*SW*_ ≥ *L_random_* and *C*_*SW*_ >> *C_random_* (Jordano *et al.*, 2003). Path lengths and clustering coefficients were calculated using weighted edges based on the estimated strength of an interaction (i.e. 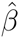); using unweighted edges does not qualitatively change the interpretation of the network.

**Figure 2:**
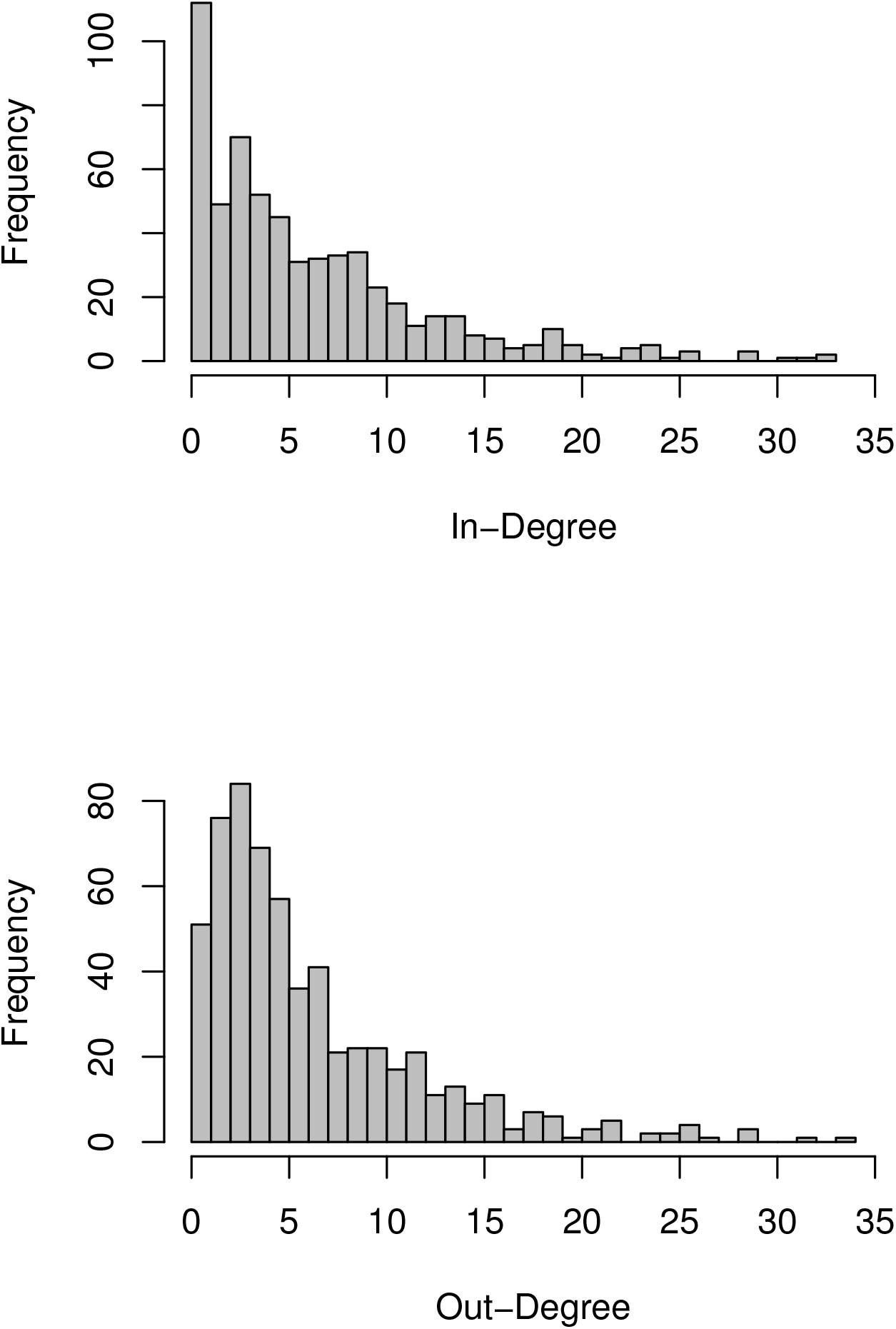
In- and out-degree distributions inferred for the woodrat gut microbiome network. In-degree is the number of OTUs that affect a focal OTU; out-degree is the number of OTUs affected by the focal OTU. A total of 4 174 interactions (edges in the network) were estimated from the AR(1) model. The network structure shows a small-world pattern that is common among many non-microbial ecological communities.

We assessed the fit of models using a pseudo-*R*^2^ metric based on the percentage of the deviance explained by the model (Cameron & Windmeijer, 1997). Figure 3 shows the distribution of pseudo-*R*^2^ values for the fitted models. Models with a high pseudo-*R*^2^ predict the dynamics of a particular OTU better than models with relatively lower pseudo-*R*^2^ (Figure 4). A broad range of values were returned that spanned all possible values (i.e. 0 to 1). Importantly, we found that the pseudo-*R*^2^ values did not depend on the *α* parameter of the elastic-net regularization, which influences the number of parameters (e.g. interactions with other OTUs) in the model (Figure 3). The dynamic patterns predicted by the model match expectations based on previous work on the effects of oxalate on the gut microbiome (Miller *et al.*, 2014, 2016). For example, Oxalobacter were predicted to increase with increasing oxalate consumption over the first 5 weeks and then decrease after test subjects were no longer fed oxalate (middle panel of Figure 4; pseudo-*R*^2^ ≈ 0.75 for the Oxalobacter model).

**Figure 3:**
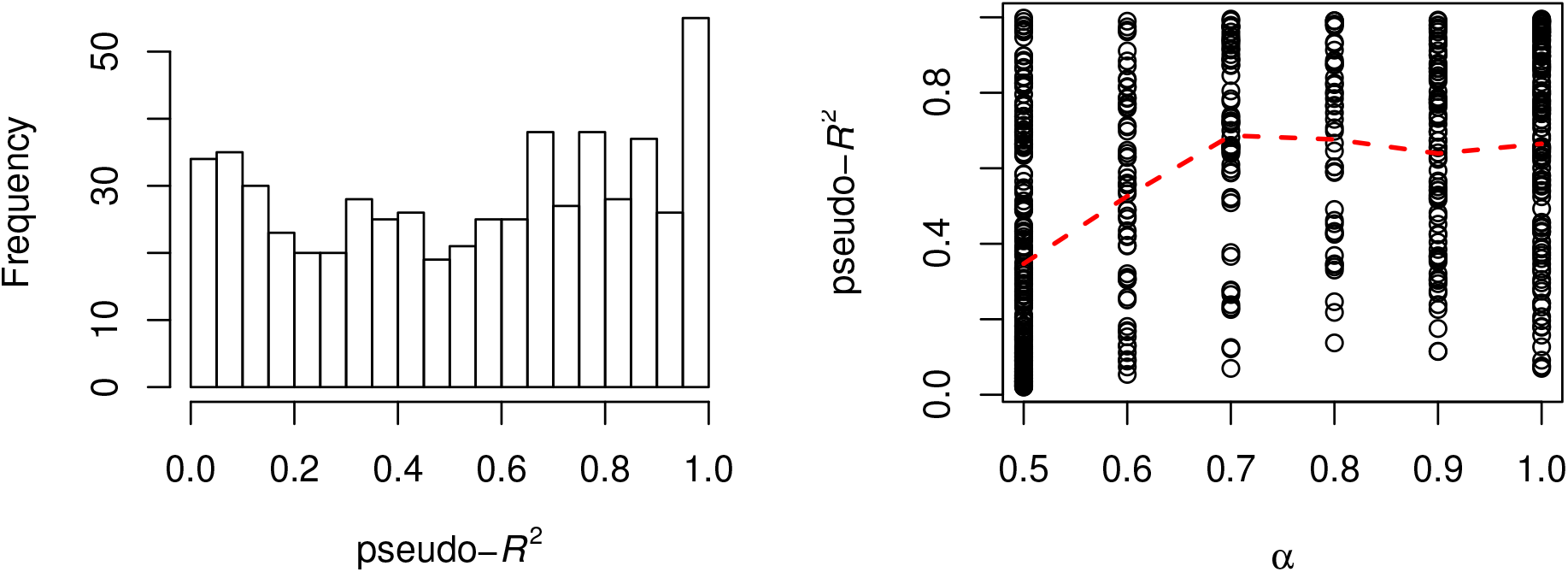
Distribution of pseudo-*R*^2^ values and their relationship to the elastic-net mixing parameter *α* from the models fit for woodrat gut microbiome data. The left panel shows that the method returned a fairly uniform distribution of values, i.e. we observed the full spectrum of poorly fitting models to very good models. The mixing parameter *α* alters the regularization penalty to favor either more small parameters (low *α*) or fewer large parameters (large *α*) in the model. The dotted red line shows the smoothed mean of the pseudo-*R*^2^ values across *α*-levels; there was no relationship between *α* (roughly, the number of parameters) and the pseudo-*R*^2^ for the AR(1) models.

**Figure 4:**
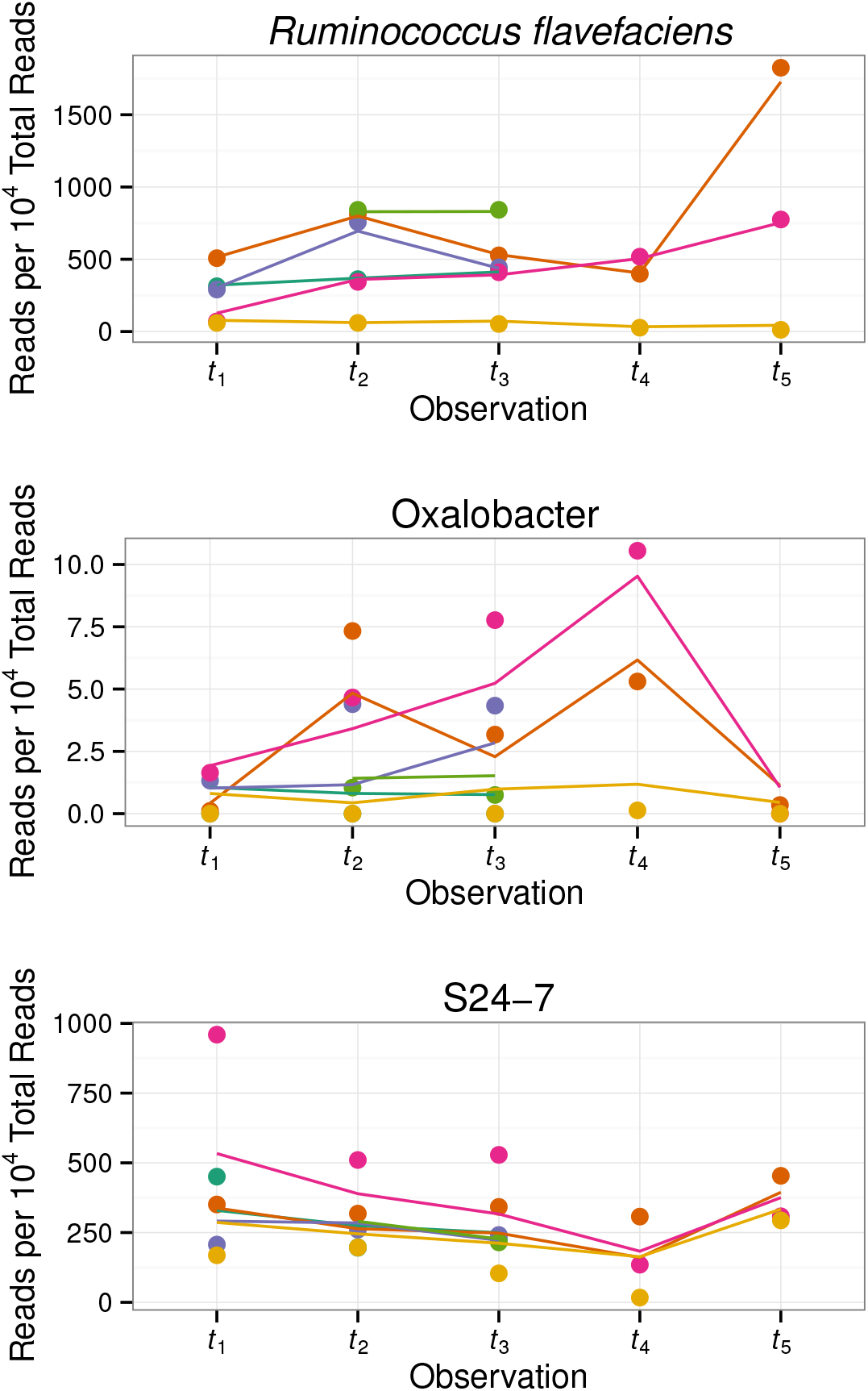
Predicted (lines) and measured (points) OTU observation rates per 10^4^ sequencing reads in the woodrat gut microbiome over the 22-day feeding the trial for three representative OTUs. Colors indicate different test subjects; some lines are incomplete because only points where data were available for the subject for *t* − 1 and *t* are plotted (i.e. those data to which an AR(1) model could be applied; this is why *t*_0_ is omitted). Note that the plots are for a single OTU, not the aggregate of all OTUs with the same taxonomic ID (e.g. the top panel is for one of 19 OTUs that was identified as *Ruminococcus flavifaciens*). The panels from upper to lower are ordered by decreasing pseudo-*R*^2^ with values of 0.99, 0.75, and 0.49, respectively. The three OTUs show different responses over time to oxalate consumption: *R. avefaciens* remains relatively constant over the entire trial; Oxalobacter increases through *t*_4_ and then decreases at *t*_5_ after oxalate consumption ceased; and S24-7 showed the opposite pattern where it decreased through *t*_4_ but rebounded in *t*_5_. See Methods for detailed observation times.

Oxalate consumption clearly has a broad range of effects on bacteria in the *N. albigula* gut (Tables 1 and 2). Figure 5 shows the distribution of these effects across all 616 OTUs, as well as the effects on oxalate excreted and digested. For the unforced model, only 8 OTUs were predicted to be affected by oxalate consumption. The results of the analysis of the woodrat data support previous findings with respect to the consumption of oxalate (Miller *et al.*, 2014, 2016). For example, we found that increased consumption of oxalate leads to increased numbers of Oxalobacter and Oxalobacteraceae within the gut (Table 2). However, we also found that other OTUs—such as some members of the Lachnospiraceae, Clostridiales, and Roseburia—are more positively affected by oxalate consumption than these well-known oxalate degraders (Table 1). The effect of oxalate consumption on these taxa may have been overlooked due to grouping OTUs together within particular taxonomic IDs. For example, while multiple OTUs within the Lachnospiraceae are strongly positively affected by oxalate consumption, several are also strongly negatively affected (Table 1); thus the *mean* effect of oxalate consumption on Lachnospiraceae is lower than that of other OTUs (Table 2) and would be overlooked in studies that aggregate OTUs.

**Figure 5:**
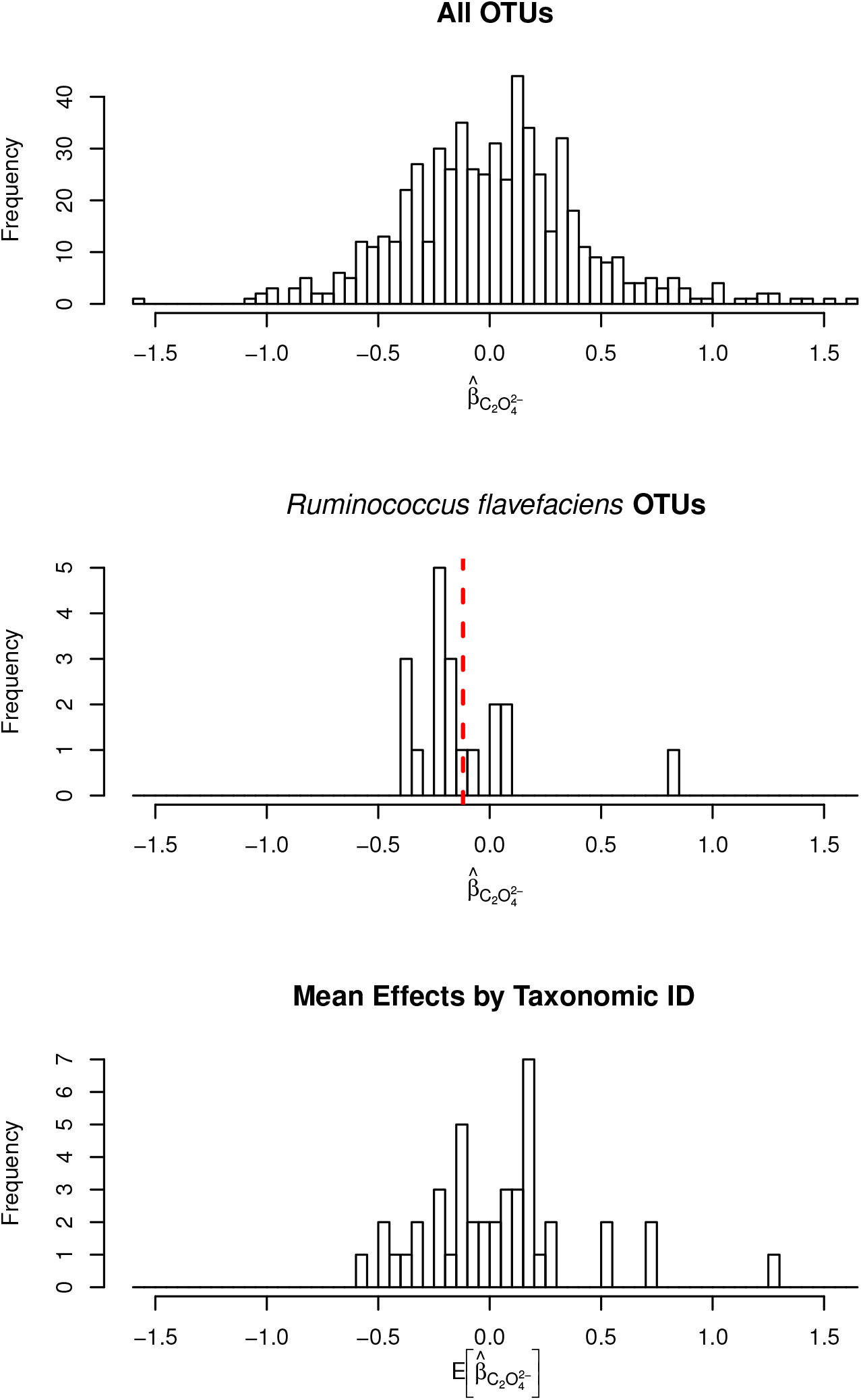
Distributions of oxalate effect sizes (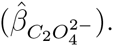 The top panel shows the distribution of effect sizes for all 616 OTUs in the woodrat gut microbiome study. The middle panel shows the distribution of estimated effect sizes for the 19 OTUs that were assigned to the species *Ruminococcus flavefaciens*; the vertical red line is the mean of those estimated effects. The bottom panel shows the distribution of mean effect sizes 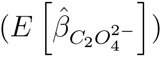 for the 43 unique taxonomic IDs to which the 616 OTUs were assigned.

**Figure 6:**
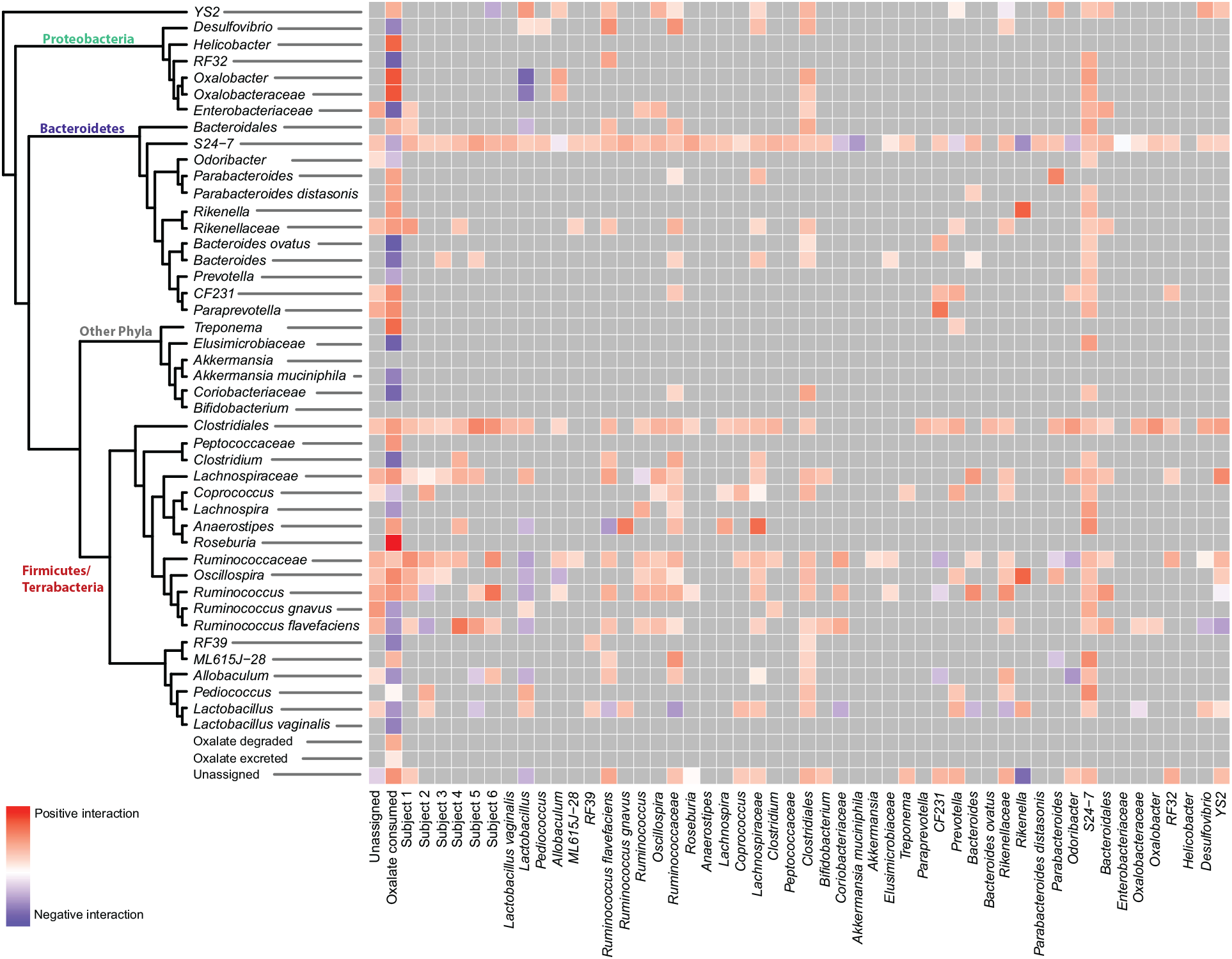
Mean pairwise effect sizes of interacting factors. The direction of the interactions is such that the factors on the horizontal axis affect the variables vertical axis (which are arranged by taxonomic classification per the Greengenes database). The scaled colors indicate the magnitude and direction of the interaction between the two variables.

**Table 1:** OTUs strongly influenced by oxalate consumption. For the forced model, the effect of oxalate consumed on the rate of observation for an OTU 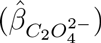 in the gut microbiome was estimated for every OTU. This table shows the top and bottom 2.5% of those estimates and the effected OTU. Note that taxonomic ID is the lowest taxonomic designation returned for a particular OTU; thus, repeats of particular IDs (e.g. Lachnospiraceae) reflect different OTUs within the particular ID.

| Negative Effects |  | Positive Effects |  |
| --- | --- | --- | --- |
| Taxonomic ID | $\hat{\beta}_{C_2O_4^{2-}}$ | Taxonomic ID | $\hat{\beta}_{C_2O_4^{2-}}$ |
| Ruminococcaceae | -1.59 | Unassigned | 1.61 |
| Lachnospiraceae | -1.06 | Lachnospiraceae | 1.54 |
| Clostridiales | -1.01 | Lachnospiraceae | 1.40 |
| Clostridiales | -1.00 | Clostridiales | 1.39 |
| S24-7 | -0.99 | Roseburia | 1.27 |
| Lachnospiraceae | -0.99 | Clostridiales | 1.25 |
| S24-7 | -0.96 | Clostridiales | 1.22 |
| Clostridiales | -0.90 | Ruminococcus | 1.20 |
| S24-7 | -0.88 | S24-7 | 1.17 |
| S24-7 | -0.87 | Clostridiales | 1.11 |
| S24-7 | -0.84 | Clostridiales | 1.04 |
| S24-7 | -0.83 | Lachnospiraceae | 1.03 |
| S24-7 | -0.83 | S24-7 | 1.02 |
| S24-7 | -0.81 | Clostridiales | 1.01 |
| S24-7 | -0.81 | Lachnospiraceae | 0.96 |

**Table 2:** Predicted effects of oxalate aggregated at different taxonomic divisions. The usual practice in microbiome research is to aggregate data at various, convenient taxonomic levels. We therefore averaged our model results across all OTUs belonging to the same taxonomic ID to get the expected effect for that group, 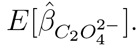 (Note that these are not nested; e.g. Oxalobacter is not part of the mean for Oxalobacteraceae.) Comparing these values to those in Table 1 shows how strain level information can be lost when aggregating OTUs.

| Negative Aggregate Effects |  |  |  | Positive Aggregate Effects |  |  |  |
| --- | --- | --- | --- | --- | --- | --- | --- |
| Taxonomic ID | $E[\hat{\beta}_{C_2O_4^2-}]$ | Division | $N_{OTU}$ | Taxonomic ID | $E[\hat{\beta}_{C_2O_4^2-}]$ | Division | $N_{OTU}$ |
| *Bacteroides ovatus* | -0.56 | species | 1 | Roseburia | 1.27 | genus | 1 |
| Elusimicrobiaceae | -0.48 | family | 1 | Oxalobacter | 0.73 | genus | 1 |
| RF32 | -0.45 | order | 4 | Oxalobacteraceae | 0.72 | family | 1 |
| Enterobacteraceae | -0.43 | family | 1 | Helicobacter | 0.55 | genus | 1 |
| Coriobacteriaceae | -0.38 | family | 1 | Treponema | 0.50 | genus | 2 |
| Clostridium | -0.34 | genus | 2 | Oscillospira | 0.27 | genus | 19 |
| Bacteroides | -0.32 | genus | 2 | CF231 | 0.25 | genus | 2 |
| RF39 | -0.22 | order | 2 | Paraprevotella | 0.24 | genus | 1 |
| *Akkermansia muciniphila* | -0.21 | species | 1 | Unassigned | 0.20 |  | 15 |
| *Lactobacillus vaginalis* | -0.21 | species | 1 | Ruminococcus | 0.19 | genus | 13 |

## Discussion

We have presented a method for the analysis of microbial community data that leverages the power of regularization techniques to infer ecological interactions based on OTU-level 16S rDNA read counts and applied these methods to time-series microbiome data from an oxalate feeding trial in woodrats. Our modeling strategy for 16S rDNA amplicon data provides a flexible and relatively computationally inexpensive method for researchers to estimate the strength of ecological interactions in microbial communities. By modeling read count data directly and using elastic-net regularization to select and stabilize the model, the method overcomes many common challenges in analyzing microbiome data.

The vast diversity of taxa that occurs in most microbiomes (Shade *et al.*, 2014; Coyte *et al.*, 2015) poses an enormous challenge in studying ecological dynamics of these systems. Computational hurdles make it intractable for many currently used methods for time-series microbiome data, such as MARSS models (Holmes *et al.*, 2012) and generalized Lotka-Volterra models (Stein *et al.*, 2013; Marino *et al.*, 2014; Buffie *et al.*, 2015), to be fit when the dynamics of many OTUs are in question. In order to make analysis tractable, researchers typically reduce the number of OTUs being studied to just a handful that are of interest (Buffie *et al.*, 2015) or to those that are most dominant in terms of their relative abundance (Vahjen *et al.*, 2011; Marino *et al.*, 2014). Both of these options provide answers that are biased *a priori*. By limiting the study to OTUs of interest, it is impossible to discover new roles for microbes within communities because “uninteresting” OTUs would never be studied. Similarly, by only analyzing the numerically dominant species, important roles of microbes whose abundance falls below the cutoff will not be investigated; it is well known from community ecology that keystone species for communities need not be numerically dominant species (Shade *et al.*, 2014). Regularization combined with appropriate mathematical models provides a technique to analyze the entirety of the data, rather than arbitrarily selecting OTUs of interest.

Typically, amplicon data are transformed to represent relative abundances within a community by dividing the number of reads by the total number of reads in the sample (Human Microbiome Project Consortium, 2012); this normalization leads to numerous statistical complications, the two most prominent being altering the correlation structure of the data and censoring of the data at some arbitrary level (Hinkle & Rayens, 1995; Egozcue *et al.*, 2003; van den Boogaart & Tolosana-Delgado, 2008; Li, 2015). Compositional data, data whose sum is forced to be one, have a different correlation structure which can mask the true nature of the interactions between species (Lin *et al.*, 2014). For example, if one OUT's relative abundance increases, it is impossible to distinguish a hypothesis of a true increase in absolute abundance from a hypothesis of a net decrease in absolute abundance of the other members of the community. Various transformations (e.g. isometric log-ratio, centered logratio) have been applied to correct for this, but, while they provide improvements, complete resolution of this forced correlation structure by transformation is unlikely (Egozcue *et al.*, 2003). Censoring in the compositional data occurs because OTUs that had zero reads in a sample get divided by variable numbers of total read counts. For example, if one sample had 100 total reads while another had 1000 total reads, then a zero from the first sample represents <0.01 compared to <0.001 for the second. Issues regarding the analysis of censored data are well documented (Hinkle & Rayens, 1995; Egozcue *et al.*, 2003; Li, 2015), and methods are available to correct these issues (Freeman & Modarres, 2002; Lin *et al.*, 2014); the methods however are often computationally expensive (e.g. bootstrapping models over imputed values) which is problematic given the already large size and complexity of the analyses, and may never fully resolve the censoring issues. Modeling the read count data directly—as suggested herein—rather than data that has been normalized to relative abundance overcomes these statistical issues.

While we have chosen to model only the linear (first-order) terms as an AR(1) model for the woodrat gut microbiome, the ARIMA model can be modified to accommodate complex dynamics (e.g. seasonality) in a system by adjusting *p*, *d*, or *q*. Higher order terms that test for complex interactions (such a trait-mediated indirect interactions) could also be included with the caveat that altering the structure of the ARIMA or the order of the predictor terms can greatly increase the number of parameters, thus exacerbating the problem that the number of possible parameters is far greater than the number of observations. As a whole, the study of microbiome dynamics needs continued advances in modeling strategies to successfully understand the eco-evolutionary complexity of microbial communities, where higher order interactions are likely to be the rule rather than the exception.

It is important to realize that, while the methods presented here can be adapted to address many questions, the quality and amount of data collected strongly influences the quality of the results. For example, Kurtz *et al.* (2015) examined the ability to recover certain synthetic network types (e.g. scale-free versus clustered networks) from different regularization algorithms and show how the ability to recover edges (interactions) in the network is dependent on the number of samples. Additionally, tuning the model and algorithm parameters (for example, see the method of cross-validation in Computational Details) has important consequences for the resulting inference. Thus, as with any statistical analysis, it is important to examine diagnostic outputs of the models to ascertain proper performance of the method.

The combination of using a (elastic-net) regularized ARIMA model with Poisson errors tackles many issues facing the analysis of microbiome time-series data and is flexible enough to be adapted to other types of analyses. For example, while we have chosen to use Poisson-distributed errors it would be easy to switch this distribution to others that are commonly used for count data, such as either the quasipoisson or negative binomial distribution, which could handle overdispersion in amplicon counts. As demonstrated in the oxalate analysis, if the experimental design is such that it is logical to force the inclusion of certain variable(s), this can be done within the regularization algorithm; the same can be said for inclusion or exclusion of an intercept term in the model. The general method of using regularization along with Poisson errors can be applied to more basic microbiome analyses as well. For example, to ask the question of which OTUs might contribute to a particular observation of interest (e.g. which OTUs in the gut microbiome are predictive of obesity), the ARIMA equations present above could be replaced by the familiar regression equation **y** = **X***β*.

As the amount of information related to the ecology and evolution of microbial communities increases, scalable methods of statistical analysis such as the method presented here will be required to make sense of data. By utilizing regularization and a model with error structure designed for count data, this method overcomes many obstacles to interpreting microbiome dynamics, providing a much needed framework to address important eco-evolutionary questions regarding microbial communities.

## Acknowledgements

We would like to thank Jodie Nicotra for help with comments and edits on the manuscript as well as members of CMCI, IBEST, and BCB at the University of Idaho for helpful discussions on the research topic. Research reported in this publication was partially supported by the National Institute Of General Medical Sciences of the National Institutes of Health under Award Numbers P20GM104420 and P30GM103324 to University of Idaho. Funding was also provided in part by NSF DEB-1342615 to M.D. The content is solely the responsibility of the authors and does not necessarily represent the official views of the sponsoring agencies.

## Author contributions statement

B.R. and C.R. conceived the modeling; A.M. and M.D. conceived the feeding experiment; A.M. performed the feeding experiment; B.R. conducted all analyses; S.B., J.W., and J.V.L. helped refine the modeling procedures and prepare the manuscript. All authors reviewed the manuscript.

